# *In vitro* host transcriptomics during *Cryptococcus neoformans*, *Cryptococcus gattii*, and *Candida albicans* infection of South African volunteers

**DOI:** 10.1101/2024.04.05.588257

**Authors:** Ronan Doyle, Shichina Kannambath, Alan Pittman, Rene Goliath, Vinod Kumar, Graeme Meintjes, James Milburn, Mihai G. Netea, Thomas S Harrison, Joseph N Jarvis, Tihana Bicanic

**Author notes:** Corresponding author: Ronan Doyle, Clinical Research Department, London School of Hygiene & Tropical Medicine. Keppel Street, London, WC1E 7HT, United Kingdom. These authors contributed equally (Ronan Doyle, Shichina Kannambath, Joseph N Jarvis, Tihana Bicanic).

## Abstract

*Cryptococcus neoformans*, *Cryptococcus gattii* and *Candida albicans* are opportunistic fungal pathogens associated with infections in immunocompromised hosts. Cryptococcal meningitis (CM) is the leading fungal cause of HIV-related deaths globally, with the majority occurring in Africa. The human immune response to *C. albicans* infection has been studied extensively in large genomics studies whereas cryptococcal infections, despite their severity, are comparatively understudied. Here we investigated the transcriptional response of immune cells after *in vitro* stimulation with *in vitro C. neoformans*, *C. gattii* and *C. albicans* infection of peripheral blood mononuclear cells (PBMCs) collected from healthy South African volunteers. We found a lower transcriptional response to cryptococcal stimuli compared to *C. albicans* and unique expression signatures from all three fungal stimuli. This work provides a starting point for further studies comparing the transcriptional signature of CM in immunocompromised patients, with the goal of identifying biomarkers of disease severity and possible novel treatment targets.

## Major article Background

*Cryptococcus neoformans* and *Cryptococcus gattii* are ubiquitous environmental organisms to which humans are exposed via inhalation. Invasive infection is rare in immunocompetent hosts, however in individuals with deficient immunity, dissemination occurs from the lungs into the central nervous system (CNS), causing cryptococcal meningitis (CM). CM is one of the leading causes of HIV-related deaths globally, with mortality ranging from 30-70%, and over 75% of CM cases occur in Africa [1].

Studies of the immune response to cryptococci show pattern recognition of cryptococcal components, such as glucuronoxylomannan (GXM) and mannoproteins (MPs), and activation of CD4+ T cells, mainly Th1, leading to activation of M1 macrophages and yeast cell engulfment [2,3], mediated via the production of cytokines IFN-_γ_, TNF-_α_, IL-6 and IL-17, amongst others [4,5]. The association of CM with advanced HIV disease, characterised by depletion of CD4+ T cells, illustrates the importance of CD4-mediated host defense [6]. Defects in Th1-type immune responses result in production of Th2-mediated alternatively activated macrophages that allow cryptococci to survive and proliferate [7–9]. Our recent genome-wide association study, which included South African participants of African descent with advanced HIV, identified 6 single nucleotide polymorphisms upstream from CSF1 associated with susceptibility to cryptococcosis [10].

*Candida albicans* is a human gut commensal associated with mucosal infection, and another opportunistic fungal pathogen in the context of advanced HIV. Compared to *Cryptococcus*, *C. albicans* is a well-studied organism, and human studies have used transcriptional profiling to understand the host immune response to infection. *C. albicans* infections activate both innate immune cells such as neutrophils and macrophages, as well as inducing Th1 and Th17-mediated immune response [11,12]. While host defence mechanisms mediated by innate immune cells, especially neutrophils, are crucial in systemic *Candida* infections, Th17-responses mainly mediate anti-*Candida* mucosal host defence. During invasion of the tissues, *C. albicans* adapts and switches to its filamentous hyphal form, allowing escape and immune evasion [13].

Unlike *C. albicans*, immunological studies of *C. neoformans* and *C. gattii* have been limited to a handful of selected cytokine targets or murine models [4–6,14–16]. Most of the human immunological studies have been performed in European populations, while very limited data are available from populations in Africa, where the burden of CM-related mortality is greatest. Despite expanded ART access in high HIV-prevalence African settings leading to reductions in the incidence of CM, mortality remain high. Further improvement of antifungal therapy is necessary to improve outcomes and is reliant on an understanding of the host response to the infection. In this study, we undertook the first whole transcriptome study of the human response to cryptococcal infection in healthy South African volunteers.

## Methods

### Healthy human volunteer cohort

Following informed consent, 25 ml of whole blood was drawn from 15 (8 female, 7 male) healthy volunteers (confirmed HIV negative) of self-identified Xhosa ethnicity from Cape Town, South Africa. Peripheral blood mononuclear cells (PBMCs) were isolated according to a standard Ficoll-Paque plus (GE healthcare) protocol, counted and adjusted to 5x10^5^ cells/ml, then cultured in RPMI 1640 media supplemented with gentamicin 10 mg/mL, L-glutamine 10 mM, pyruvate 10 mM and 10% human serum (Sigma, UK).

### Ethical approval

The study was approved by the University of Cape Town Human Research Ethics Committee in 2014 (Ref 721/2014), as an extension of the original human GWAS study (Ref 018/2005), published separately [10].

### PBMC stimulations

5x10^5^ PBMCs were stimulated for 24h at 37^0^C with 3 fungal stimuli: *Cryptococcus neoformans* H99, *Cryptococcus gattii* R265 and *Candida albicans* UC280 (all heat-killed by incubation at 65^0^C for 2h), or remained unstimulated. The cryptococcal stimuli were opsonised with monoclonal anti-capsule (18B7) antibody (kindly provided by A Casadevall, Albert Einstein College of Medicine, New York). Cells were harvested for RNA extraction at 6h and 24h from unstimulated and stimulated PBMCs using TRIzol® (Life technologies) reagent according to manufacturer’s instructions and treated with DNase to ensure elimination of genomic DNA. RNA concentration and quality was analysed using a Qubit fluorometer (Life Technologies) and 2200 Tape station (Agilent Technologies), respectively.

### RNA sequencing and analysis

RNA sequencing libraries were prepared using the TruSeq RNA Library Preparation Kit v2 (Illumina), according to manufacturer’s instructions. Libraries were then sequenced to generate 150-bp paired-end reads on an Illumina Novaseq 6000 using an S4 flow cell. Read quality was assessed using FASTQC (https://www.bioinformatics.babraham.ac.uk/projects/fastqc/) and samples with too few reads prior to alignment were removed using fastp software (version 0.20.0) (https://github.com/OpenGene/fastp). Sequences were aligned to the human reference genome (hg38) with STAR (version 2.6.8a) [17] and samples with a low percentage of reads mapped to the reference were removed. Differential gene expression was performed using DESeq2 (v1.30.1) [18] in R (v3.8.3). Gene ontology analysis was carried out using goseq (v1.54.0) [19], also in R. Pathway analysis was carried out using Reactome [20].

### Data availability

The raw RNA sequencing data produced during this study is split over two repositories. It is available from figshare via https://figshare.com/s/b953f3192c77cef0be98 and the European Nucleotide Archive under study accession number: PRJEB74561.

## Results

### Cryptococcal-induced Differential Gene Expression in Human PBMCs

To understand the transcriptional response to cryptococcal stimulation, we compared PBMCs from 15 healthy donors stimulated with two cryptococcal species commonly causing human CM (*Cryptococcus neoformans* and *Cryptococcus gattii*) with unstimulated cells. RNA was isolated from PBMCs at 6h and 24h post stimulation, andmRNA was then sequenced from each sample, generating a mean of 36.6 million reads per sample. We determined the differentially expressed genes (DEGs) at both time points using a cut-off of >1 log2 fold change in expression and Benjamini and Hochberg’s (BH)-adjusted *P* value < 0.05. *C. gattii* had the greater number of DEGs at 6h post-stimulation (280 vs 30), whereas *C. neoformans* had more abundant DEGs at 24h post-stimulation (71 vs 17) (Figure 1). There were 5 upregulated and 25 downregulated DEGs for *C. neoformans* at 6h, and 3 upregulated and 68 downregulated DEGs after 24h. For *C. gattii* we found 4 upregulated and 276 downregulated DEGs at 6h and 0 upregulated and 17 downregulated DEGs after 24h. The full list of DEGs associated with *C. neoformans* and *C. gattii* stimulation at both time points can be found in supplementary table 1.

**Figure 1.**
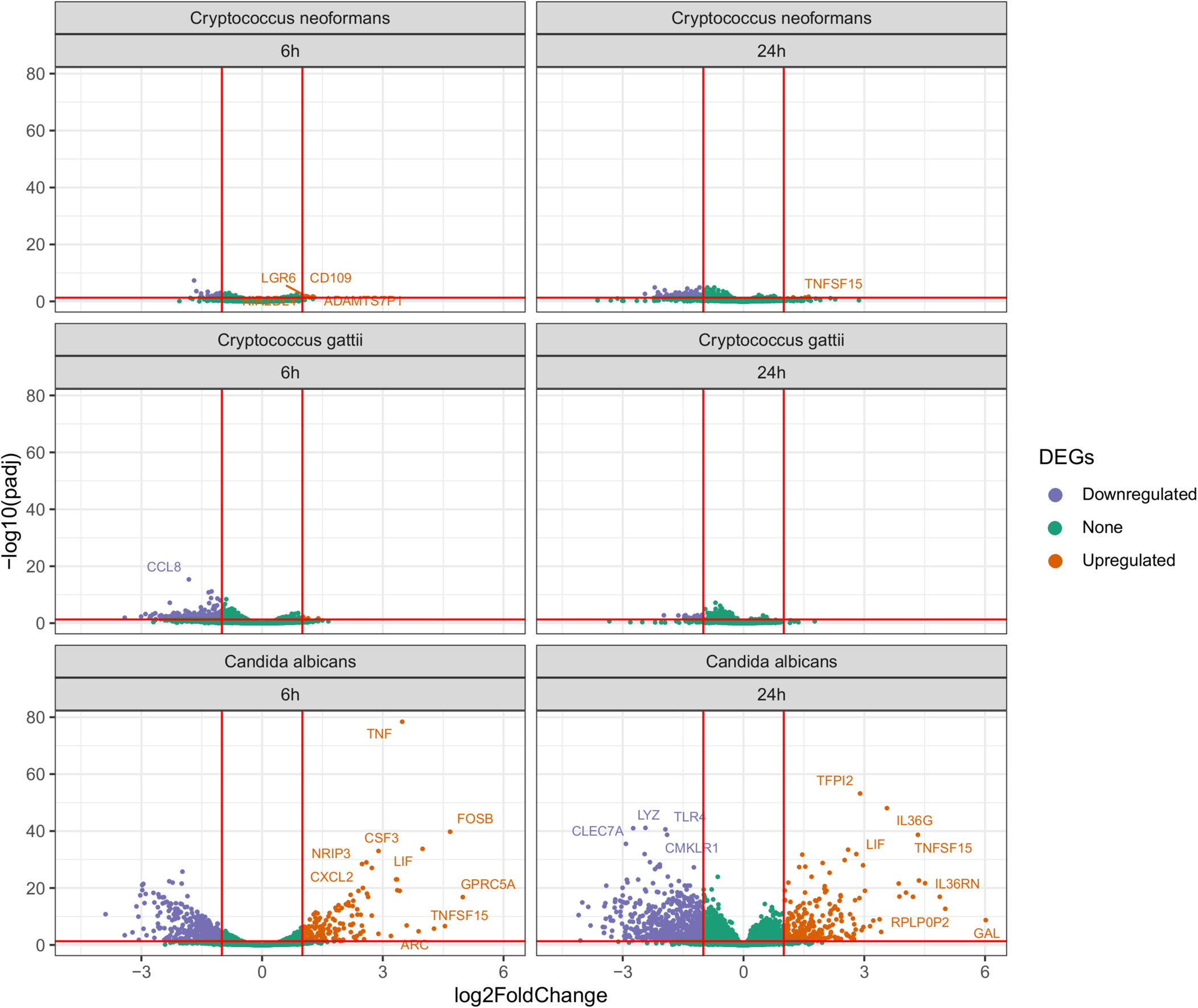
Volcano plots showing the number of differentially expressed genes when compared to unstimulated controls for C. neoformans, C. gattii and C. albicans at 6h and at 24h.

We compared the fraction of shared DEGs between the two time points for both pathogens and found little (< 1%) to no crossover between gene expression within species at the two time points (Figure 2). *C. neoformans* shared no DEGs between 6 and 24h time points, whereas *C. gattii* shared 2 DEGs between 6 and 24h. One of which one was an RNA pseudogene (*RN7SL4P)* and the other was a long intergenic non-protein coding RNA (lncRNA) gene (*LINC01094)*.

**Figure 2.**
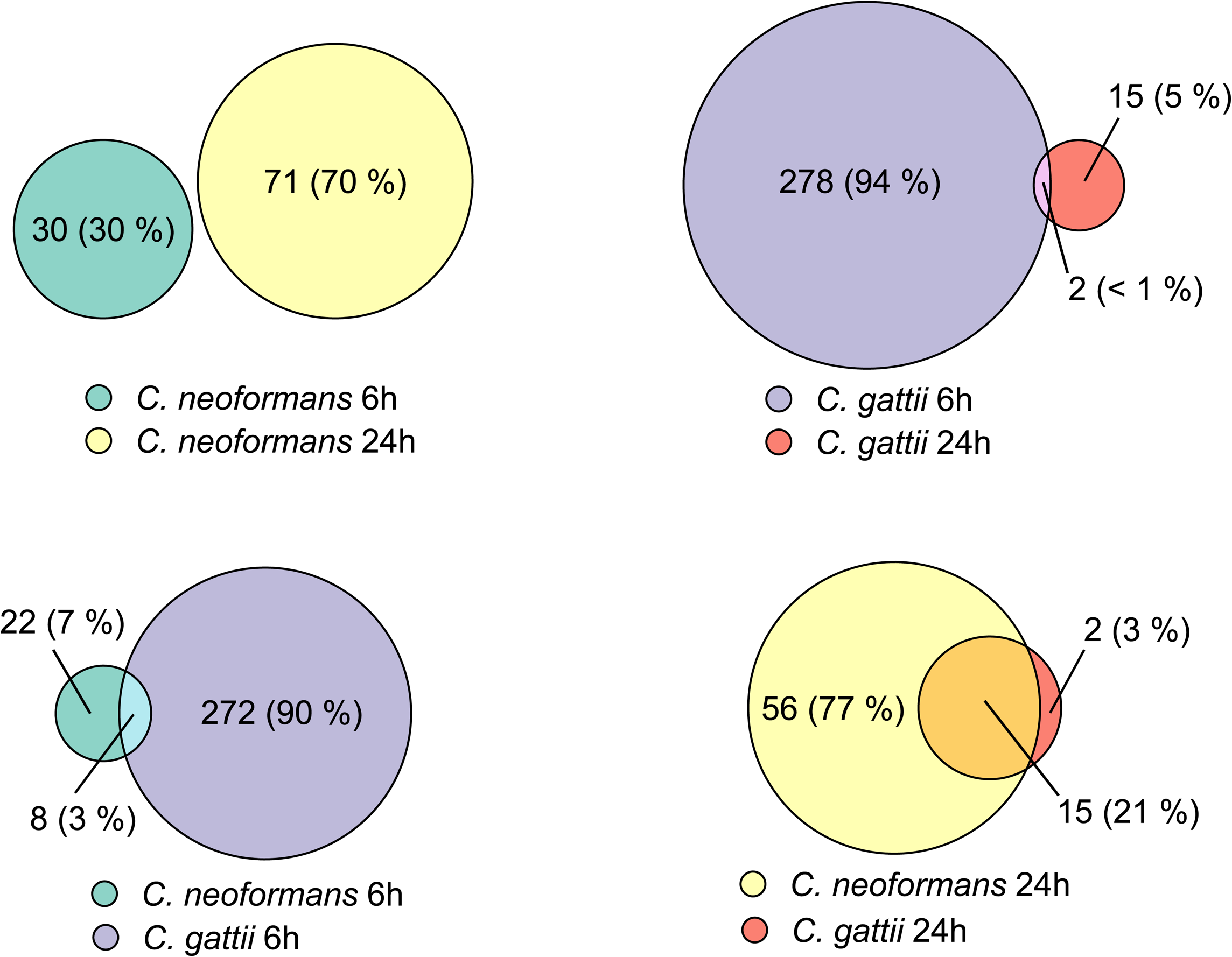
Euler diagram showing number of differentially expressed genes shared between C. neoformans at two different time points. Then the same comparison between C. neoformans and C. gattii at the same time points.

There were also only a minority of shared DEGs between species at the same time points; 3% (8 genes) of total DEGs were shared at 6h and this rose to 21% (15 genes) of DEGs at 24h.

### Cryptococcal Pathway Analysis

We used Gene Ontology (GO) terms to functionally annotate these DEGs found post-stimulation with *C. neoformans* and *C. gattii*. Analysing the GO molecular function pathways at 6h post-stimulation showed *C. neoformans* had 7 pathways significantly altered compared to unstimulated controls, whereas *C. gattii* had 13 (Figure 3). Both pathogens had significantly enriched pro-inflammatory pathways containing genes associated with cytokine and chemokine activity and receptor binding. GO molecular function pathways significantly differentially expressed in *C. gattii* at 6h post-stimulation included Toll-like receptors and CXCR3 chemokine receptors. GO biological processes significantly associated with *C. gattii* included neutrophil chemotaxis and genes associated with a cell defence response to fungi. GO biological processes significantly associated with *C. neoformans* 6h post-stimulation included pro-inflammatory pathways associated with T-cell activation and differentiation (Supplementary Figure 1). By 24h, none of these 6h pro-inflammatory pathways remained significantly enriched for either cryptococcal stimulus.

**Figure 3.**
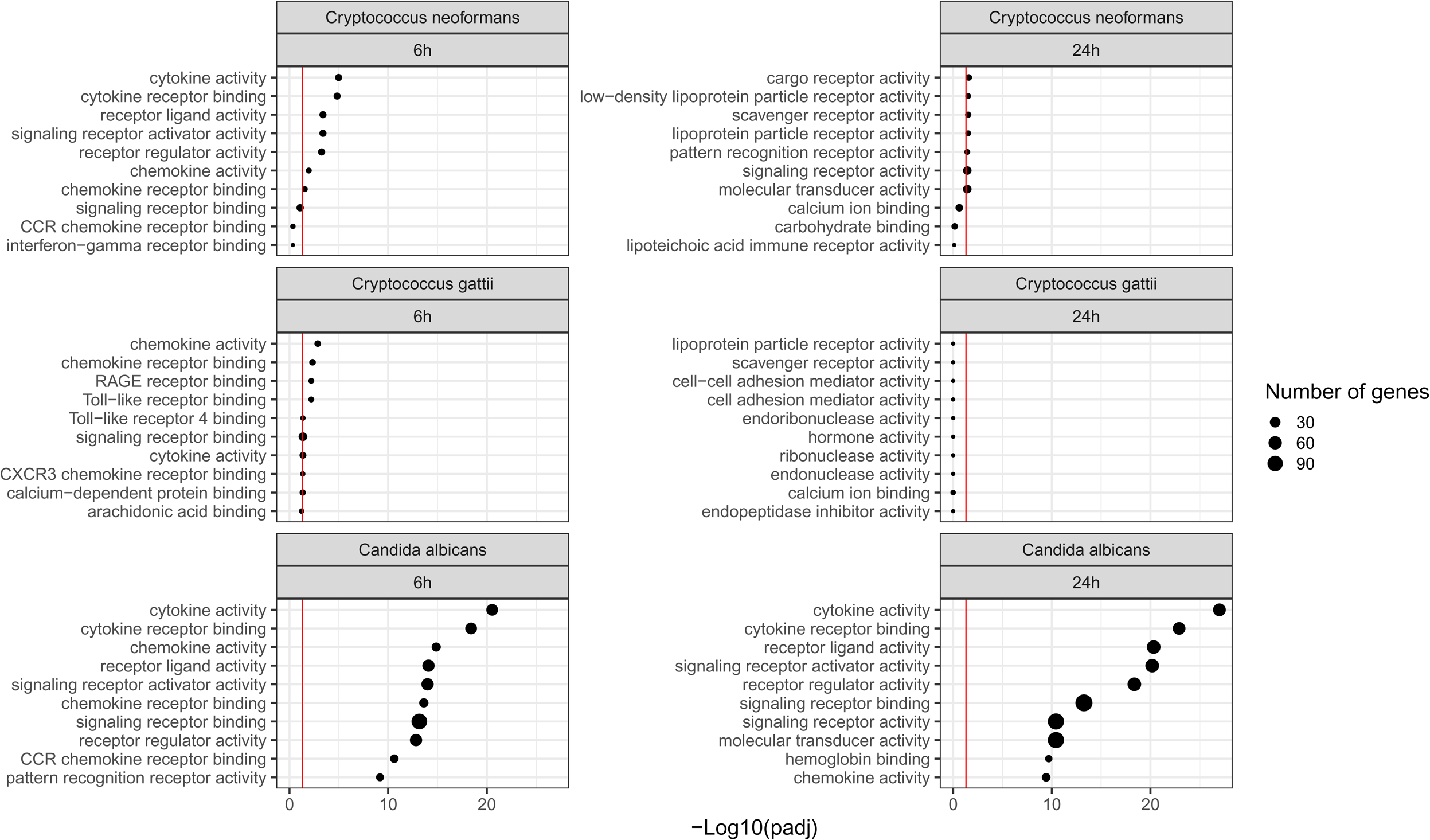
Gene Ontology molecular function pathways enriched in Cryptococcus neoformans, Cryptococcus gattii and Candida albicans at 6h and 24h post-stimulation. Top 10 pathways displayed by lowest adjusted P-value and red line denotes adjusted P value<0.05.

### *Candida albicans*-induced Differential Gene Expression in Human PBMCs

We compared cryptococcal responses to *Candida albicans* as a model for a previously well-characterised anti-fungal immune response. As described previously, we stimulated PBMCs from the same 15 healthy donors with heat-killed *C. albicans* and isolated RNA at 6h and 24h. When compared to the heat-killed cryptococcal species, a much more pronounced inflammatory response was seen in *C. albicans* stimulated cells at the same time points. There were 580 DEGs at 6h post-stimulation with *C. albicans* and 927 at 24h, compared to 280 and 30 at 6h, and 17 and 71 at 24h for *C. gattii* and *C. neoformans,* respectively. For *C. albicans* DEGs this corresponded to 413 downregulated and 167 upregulated at 6h and 656 upregulated and 271 downregulated at 24h. The full list of DEGs associated with *C. albicans* stimulation at both time points can be found in supplementary table 1.

When we compared the fraction of shared DEGs for *C. albicans* between 6h and 24h, we found 26% (Figure 4) of the total number of DEGs were shared between both time points, compared to the completely distinct cryptococcal responses at 6h and 24h. At 24h post-stimulation, almost the entirety of the transcriptional response found in *C. neoformans* and *C. gattii* stimulated cells matched with the DEGs found in *C. albicans* stimulated cells (70/73 DEGs or 96%). However, as noted above, the overall scale of the response from exposure to *C. albicans* was greater than that of the cryptococcal stimuli (Figure 1).

**Figure 4.**
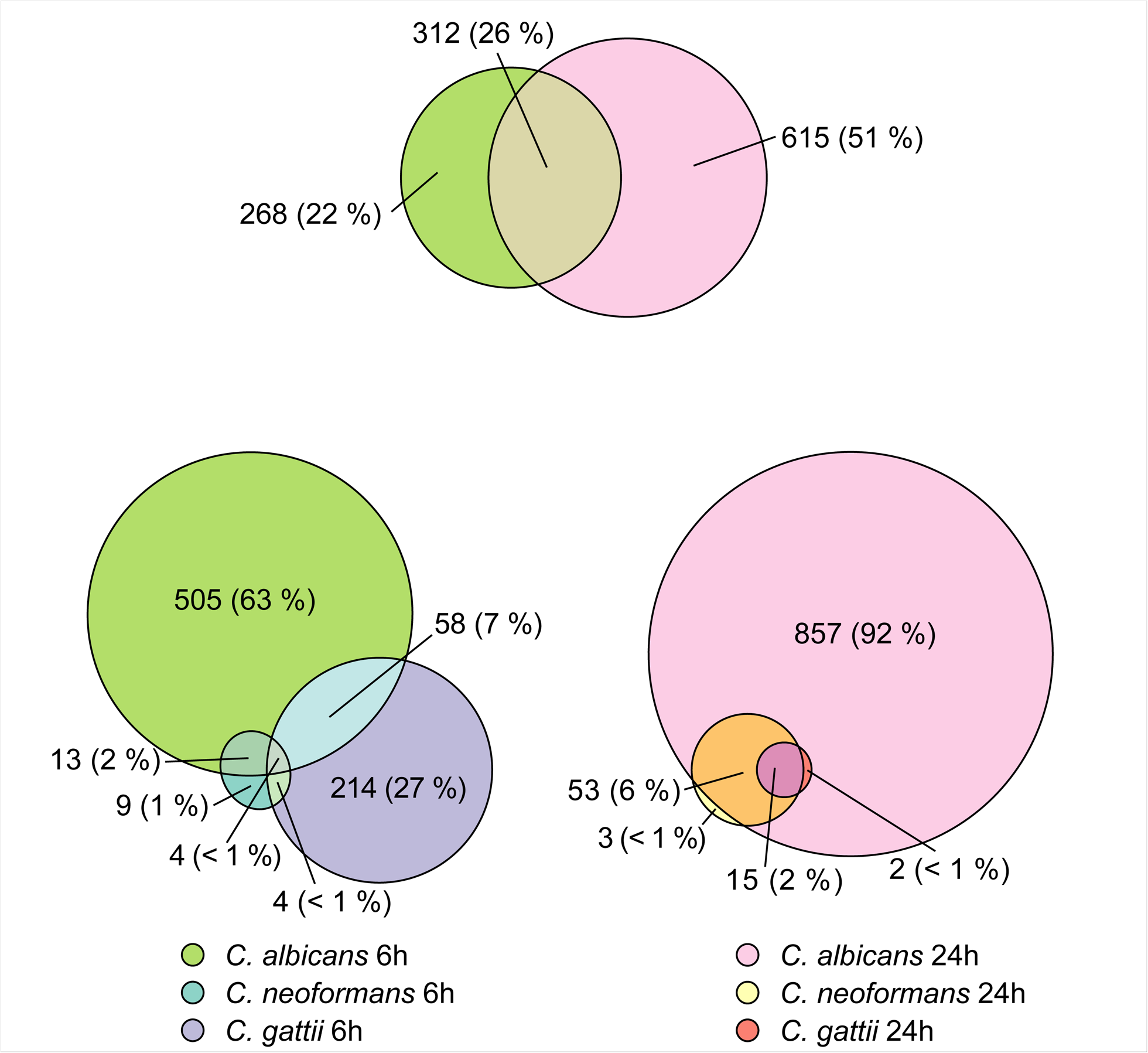
Euler diagram showing number of differentially expressed genes shared between C. albicans at two different time points. Then the same comparison between C. neoformans, C. gattii and C. albicans at both 6h and 24h post-stimulation.

As with the cryptococcal stimuli, we used GO terms to functionally annotate the DEGs found post-stimulation with *C. albicans* (Figure 3). Similar to *C. neoformans* and *C. gattii*, pro-inflammatory genes within cytokine, chemokine and receptor binding pathways were significantly enriched. However, the scale of enriched pathways was greater than the other cryptococcal stimuli and included additional pathways such as a CCR chemokine receptors and IgG binding. Comparing GO biological process pathways, at 6h, among the pro-inflammatory pathways significantly enriched were those associated with myeloid leukocyte migration, whereas by 24h this was replaced by cytokine production and its regulation.

### Comparative host transcriptomics between cryptococcal and C. albicans stimulations

We further wanted to determine whether there were unique features of cryptococcal inflammatory responses compared to those following *Candida* stimulation, as well as any differences between *C. neoformans* and *C. gattii*. As previously noted, no inflammatory pathways were significantly altered at 24h post-cryptococcal stimulation. We therefore focused on the 6h post-stimulation timepoint, as this comprised pro-inflammatory signatures common to all three species (Figure 3 & 5). There were 505 unique DEGs associated with *C. albicans* stimulation, 214 associated with *C. gattii* stimulation and only 6 unique DEGs associated with *C. neoformans* stimulation (figure 3). To determine the role of this group of genes, we again functionally annotated them using GO terms as above, retaining only genes which had a higher order GO Biological process annotation corresponding to “immune system process”. We used Reactome pathway analysis to group these genes into specific inflammatory pathways (Figure 5). We found only a single pro-inflammatory DEG associated with all 3 pathogens, CXCL-10, a chemokine strongly induced by interferon-gamma and recognised as a component of the Il-10 signalling pathway in Reactome pathway database. *C. albicans* and *C. gattii* shared the highest number of inflammatory DEGs (15 DEGs) compared to *C. neoformans* and *C. albicans* (4 DEGs) and we found no DEGs shared solely between *C. neoformans* and *C. gattii*. Common pathways shared between *C. albicans* and *C. gattii* included Dectin-2 C-type lectin receptors and the complement cascade, all of which are important elements of the innate immune response to these fungi. Pathways enriched uniquely following *C. neoformans* stimulation included those associated with Killer Ig-Like Receptors (KIRs) and Regulatory T cells (Tregs).

**Figure 5.**
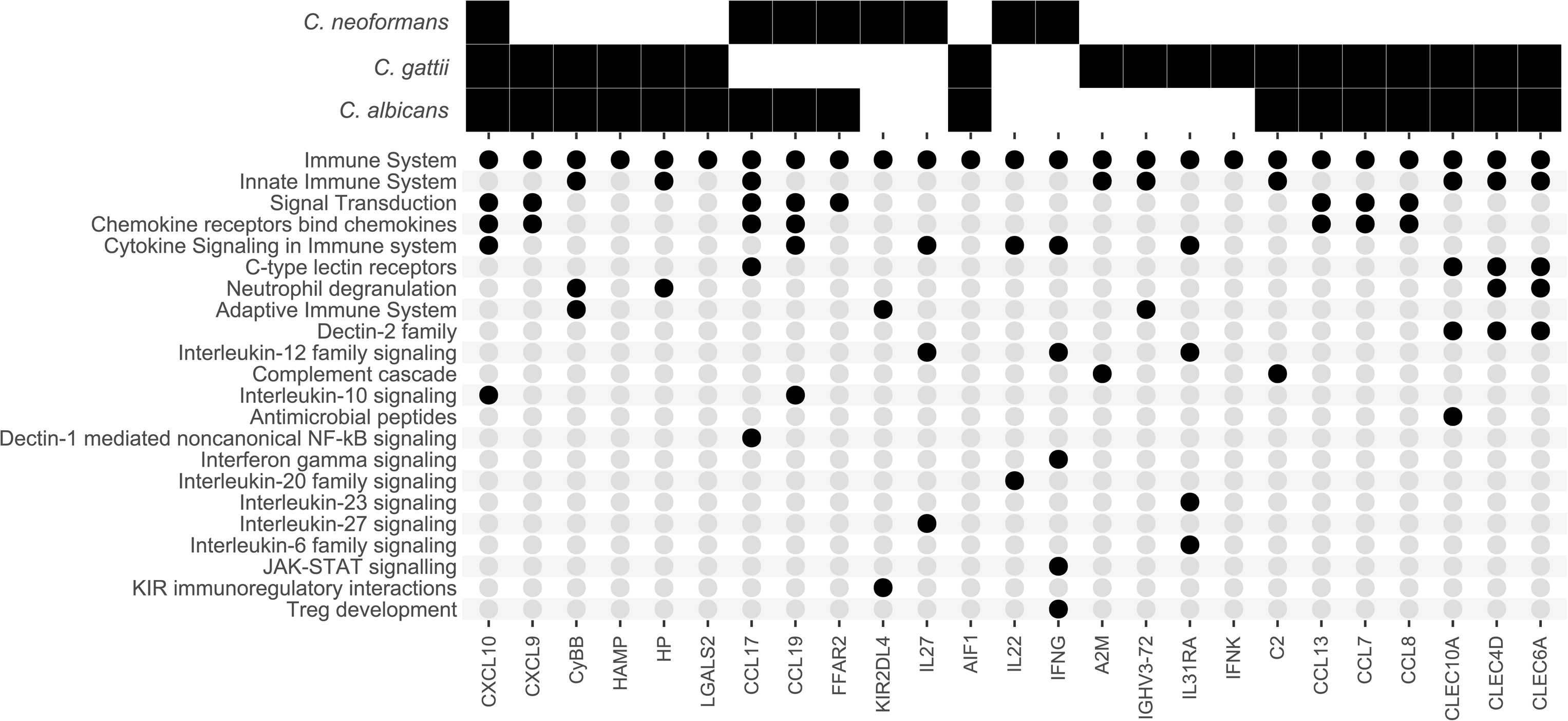
Expression of pro-inflammatory genes following heat-killed C. neoformans, C. gattii and C. albicans stimulation. A solid black square denotes a gene that is significantly differentially expressed in specific pathogen samples when compared to unstimulated controls. Black circles correspond to the Reactome immunological pathways each gene is associated with.

## Discussion

In this study we describe for the first time the transcriptome response to stimulation of human PBMCs by commensal and environmental fungal pathogens in healthy South African volunteers, and we show unique gene expression signatures in *C. neoformans*, *C. gattii* and *C. albicans*. The scale of the transcriptional response following stimulation by both cryptococcal species was considerably smaller than that to *C. albicans*. The cryptococcal inflammatory response peaked at 6h and had subsided by 24h, whereas the *C. albicans* response persisted at 24h post-stimulation.

There are many possible reasons for these differences. *C. albicans* is a human commensal of the gastrointestinal and vulvovaginal tracts causing mucosal infection in advanced HIV disease and invasive disease (candidiasis) in hospitalised patients, often in the context of mucosal barrier breach [21], whilst *Cryptococcus* is an environmental saprophyte acquired by inhalation, causing invasive infection (CM) largely in immunocompromised hosts. Human immune responses to *Cryptococcus* are triggered by phagocytosis of encapsulated cryptococcal cells (enhanced by opsonising antibodies and complement) by macrophages/ dendritic cells and subsequent antigen presentation to T cells (CD4+, CD8+ and NK cells), orchestrated and potentiated by pro-inflammatory cytokines such as TNF_α_ and IFN-_γ_. The capsule is anti-phagocytic and disrupts T cell proliferation, which could explain the paucity of inflammatory responses seen in our study. Of the two cryptococcal pathogens, *C. gattii* is responsible for relatively more disease in immunocompetent hosts [22,23] and this seems to corroborate with the more pronounced inflammatory response when compared to *C. neoformans*, as previously shown.

There has not been a prior study of the human transcriptome response to fungal pathogens from immunocompetent African volunteers. A study in China found similar upregulated pathways, such as NF-kB signalling and JAK-STAT pathway signalling, following heat-killed *C. neoformans* stimulation of 15 healthy volunteers [24]. We found little overlap in the individual DEGs between these two studies, which may underline differences in either host or pathogen-related factors, as well as methodological differences (the Chinese study used monocytes alone and a different timepoint of 12 hours).

The inflammatory response to *C. albicans* was of a greater magnitude compared to *C. neoformans* and *C. gattii* and included some overlap in DEGs. However, there was only a single inflammatory Reactome pathway over-represented in all 3 fungal species, which was IL-10 signalling through the differential expression of CXCL10. This has previously been identified as an protective or at least secondary response to regulate inflammation in all 3 pathogens [5,11,25,26]. Other than this pathway, the inflammatory response caused by heat-killed *C. neoformans* and *C. gattii* seem to be distinct from each other. A transcriptomics study that infected mice using both *C. neoformans a*nd *C. gattii*, also identified distinct gene expression signatures caused by the two fungal species, but they were not entirely unique [27]. Although there were differences, the study also found classical complement activation as unique to *C. gattii* infection, compared to *C. neoformans*, and expression of natural killer cell genes, such as receptor *KIR2DL4*, unique to *C. neoformans*.

Another significantly expressed gene unique to *C. neoformans* stimulation was interferon-gamma, which we and others have previously described as key in the clearance of infection [4,28]. IL-27 was also significantly differentially expressed in *C. neoformans* samples, and has been associated with the well-characterised cryptococcal immune reconstitution inflammatory syndrome [28].

A feature common to all the fungal stimuli includes innate immune recognition of fungal cell wall antigens. We identified shared pathways between *C. neoformans* and *C. albicans* that included Dectin-1 mediated NF-kB signalling; between *C. gattii* and *C. albicans* we identified enhanced expression of genes encoding for Dectin-2 family signalling. For *C. albicans*, important innate immune pattern recognition receptors are C-type lectin receptors (CLR) of the Dectin-1 cluster which recognise the (1,3)-Beta-d-glucans (BDG) in its cell wall [29]. Dectin-2 is another CLR that recognises the alpha-mannan constituent of the *C. albicans* cell wall [30] and has also been previously shown to bind the same proteins in the *C. gattii* cell wall [31].

Limitations of this study include the relatively small number of healthy controls and a lack of comparison to a diseased cohort. Further studies should perform RNA sequencing from immune cells from blood or CSF collected from patients with confirmed cryptococcal meningitis to identify differential gene expression associated with mortality and fungal clearance. Additional comparisons should include other pathogens common to this specific patient population, such as *Mycobacterium tuberculosis*.

In conclusion, this study has used de novo transcriptomic sequencing to identify unique gene expression signatures that differentiate *C. neoformans* and *C. gattii* infection from each other, and from *C. albicans* infection, in a healthy African control population. This work provides a foundation for further transcriptomic studies to investigate the gene expression signature associated with mortality in cryptococcal meningitis and provide new insights into potential novel treatment targets.

## Acknowledgements

This work was funded by the National Institute for Health Research (NIHR) through a Global Health Research Professorship to JNJ (RP-2017–08-ST2-012) using UK aid from the UK Government to support global health research, and by a Wellcome Trust Strategic Award for Medical Mycology and Fungal Immunology 097377/Z/11/Z, via a postdoctoral fellowship award to TB and MN. The views expressed in this publication are those of the authors and not necessarily those of the NIHR or the UK Department of Health and Social Care.

We thank the volunteers who participated in the study, nurses at Site B Khayelitsha who organised recruitment and consent for the study; Professor Robert Wilkinson, University of Cape Town, in whose laboratory the PBMC stimulation and RNA extractions were conducted.

