## Supplementary figures and images for "*In vitro* host transcriptomics during *Cryptococcus neoformans*, *Cryptococcus gattii*, and *Candida albicans* infection of South African volunteers"

### Supplemental Figure 1

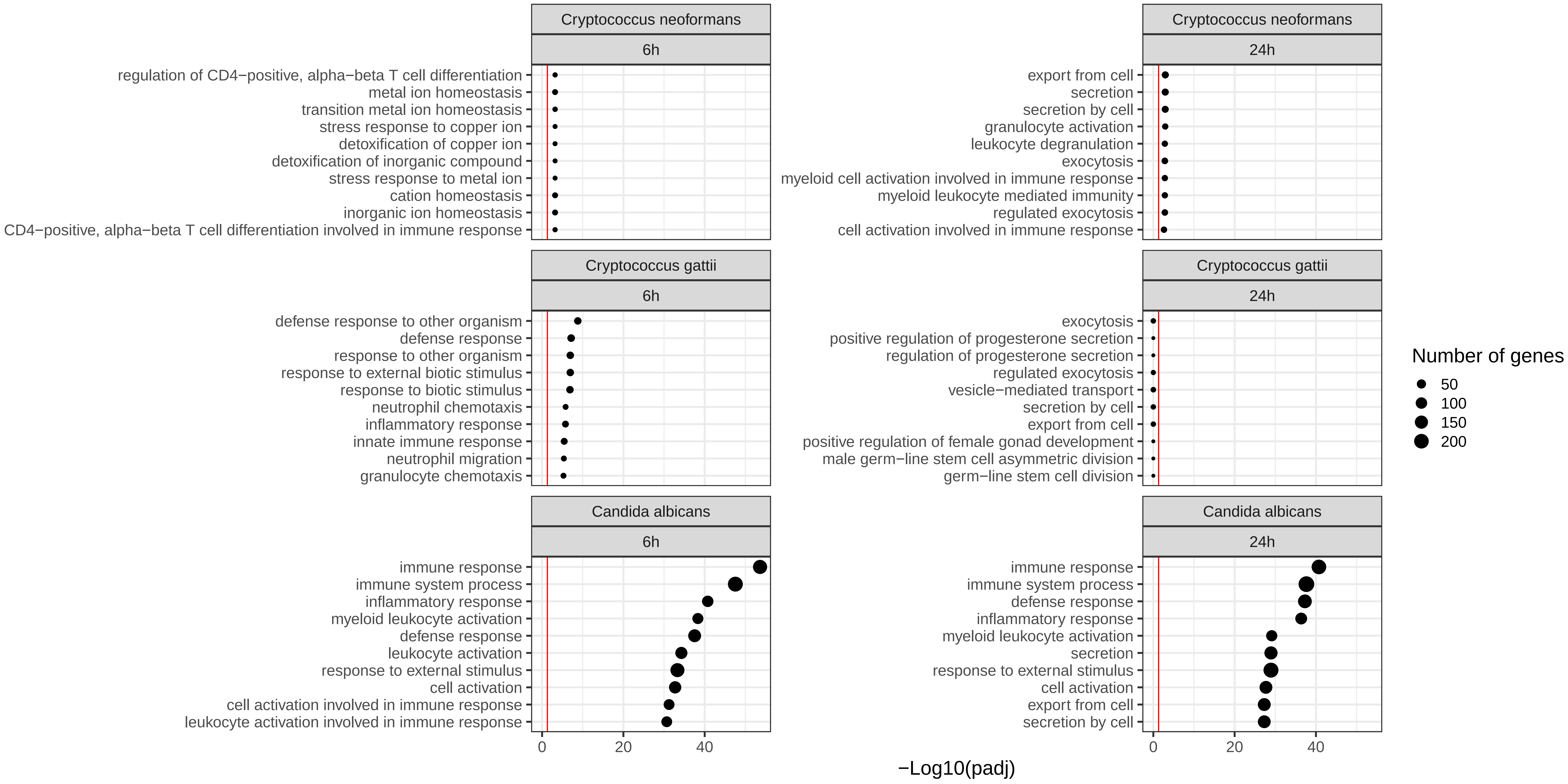
